# Excess dietary sodium partially restores salt and water homeostasis caused by loss of the endoplasmic reticulum molecular chaperone, GRP170, in the mouse nephron

**DOI:** 10.1101/2024.01.13.575426

**Authors:** Aidan Porter, Hannah E. Vorndran, Allison Marciszyn, Stephanie M. Mutchler, Arohan R. Subramanya, Thomas R. Kleyman, Linda M. Hendershot, Jeffrey L. Brodsky, Teresa M. Buck

## Abstract

The maintenance of fluid and electrolyte homeostasis by the kidney requires proper folding and trafficking of ion channels and transporters in kidney epithelia. Each of these processes requires a specific subset of a diverse class of proteins termed molecular chaperones. One such chaperone is GRP170, which is an Hsp70-like, endoplasmic reticulum (ER)-localized chaperone that plays roles in protein quality control and protein folding in the ER. We previously determined that loss of GRP170 in the mouse nephron leads to hypovolemia, electrolyte imbalance, and rapid weight loss. In addition, GRP170-deficient mice develop an AKI-like phenotype, typified by tubular injury, elevation of clinical kidney injury markers, and induction of the unfolded protein response (UPR). By using an inducible GRP170 knockout cellular model, we confirmed that GRP170 depletion induces the UPR, triggers an apoptotic response, and disrupts protein homeostasis. Based on these data, we hypothesized that UPR induction underlies hyponatremia and volume depletion in rodents, but that these and other phenotypes might be rectified by supplementation with high salt. To test this hypothesis, control and GRP170 tubule-specific knockout mice were provided with a diet containing 8% sodium chloride. We discovered that sodium supplementation improved electrolyte imbalance and reduced clinical kidney injury markers, but was unable to restore weight or tubule integrity. These results are consistent with UPR induction contributing to the kidney injury phenotype in the nephron-specific GR170 knockout model, and that the role of GRP170 in kidney epithelia is essential to both maintain electrolyte balance and cellular protein homeostasis.

## INTRODUCTION

The kidney is indispensable for electrolyte and water homeostasis, blood pressure control, and waste excretion. Acute kidney injury (AKI), the rapid deterioration of renal function, is marked by rising serum creatinine, oliguria, toxin accumulation, and disruption of water and electrolyte homeostasis. Affecting ∼15% of hospitalized patients, AKI significantly increases in-hospital mortality, and contributes to ∼2 million deaths annually **(1)**. AKI has also been linked to hypertension, chronic kidney disease, and the need for future dialysis **(2)**. The kidney, which receives about 25% of cardiac output, is especially vulnerable to injury because of its metabolically-taxing responsibilities, as well as the osmolar, hemodynamic, and toxic stressors to which nephrons are exposed **(3)**.

Each nephron segment is lined by specialized epithelial cells that express myriad transmembrane channels and transporters **(4)**. These epithelial cells in the proximal tubule (PT) reabsorb the vast majority of the glomerular filtrate, including ∼70% of sodium, and virtually all urinary protein, amino acids, and glucose **(5–7)**. The PT is also essential for activating Vitamin D, gluconeogenesis, renin production, solute secretion, and generating ammonia **(7–12)**. These energy-intensive processes likely explain the abundance of mitochondria in the PT epithelium—second only to the density in the myocardium—and the importance of the PT in AKI pathogenesis **(13, 14)**. Downstream of the PT, the distal convoluted tubule (DCT) and collecting duct, which are under the control of aldosterone and antidiuretic hormone, are vital for fine-tuning sodium, potassium, and water reabsorption largely via the epithelial sodium channel (ENaC), the renal outer medullary potassium channel (ROMK), aquaporin 2 (AQP2), and aquaporin 4 (AQP4) **(15–17)**.

Because of the crucial role of transmembrane channels and transporters in renal physiology, the activities associated with endoplasmic reticulum (ER)—which oversees the biogenesis of nearly all membrane and secreted proteins—are indispensable for kidney function and, accordingly, the health of renal and all other cells in the body. To support ER function, molecular chaperones within the ER lumen facilitate protein synthesis, maturation, and quality control **(18–20)**. For example, our prior studies established a direct role for one such ER lumenal chaperone, GRP170 (the product of the *HYOU1* gene), during the biogenesis of the epithelial sodium channel, ENaC. Specifically, we showed that GRP170 facilitates the proteosome-dependent ER-associated degradation (ERAD) of unassembled αENaC subunits in yeast, mammalian cells, and *Xenopus oocytes* **(21–23)**.

GRP170 is an Hsp70-like protein and member of the large Hsp70 family that includes the cytosolic Hsp110 chaperones**(24)**. GRP170 also contributes to the biogenesis and/or quality control of mutant forms of α1-antitrypsin and insulin, and is linked to tumor survival and cholera toxin pathogenicity **(25–30)**. In addition to its ability to bind and maintain the solubility of non-native proteins, GRP170 is also a nucleotide exchange factor for the ER Hsp70 homolog and ER stress sensor, BiP (GRP78). GRP170 function has additionally been implicated in kidney function and a variety of other conditions arising from the accumulation of immature proteins within the ER **(31–33)**.

To better delineate the relationship between ER molecular chaperones, ER protein homeostasis (proteostasis), and renal physiology, we generated a doxycycline (dox)-inducible, nephron-tubule-specific conditional GRP170 knockout (KO) mouse (GRP170^NT-/-^) (20). GRP170 deletion in renal tubular epithelial cells compromised nephron function, leading to inappropriate naturesis, hyponatremia, hyperkalemia, impaired urinary concentrating ability, weight loss, and hyperaldosteronism. The phenotype of GRP170^NT-/-^ mice resembled that of mice lacking or deficient in the α, β or γ ENaC subunits (34–37), but in contrast to these models, GRP170^NT-/-^ mice also exhibited a profound AKI-like phenotype: elevated serum BUN and creatinine, and NGAL and Kim1 expression. In addition, the GRP170^NT-/-^ animals developed albuminuria and glucosuria, hallmarks of PT injury, and renal histology revealed features consistent with PT injury. As predicted based on our prior work, the loss of GRP170 also altered nephron channel stability and trafficking, especially in the proximal nephron(22, 23, 38).

As noted above, ER homeostasis is critical for cellular health, but when ER function is blunted—by for example an increase in misfolded proteins or cellular stressors such as oxidative injury or ischemia—a stress response known as the unfolded protein response (UPR) is induced. The UPR has been linked to numerous kidney diseases, including AKI (39, 40). In response to stress, UPR activation is initially an adaptive mechanism that restores ER proteostasis by upregulating the production of ER chaperones, including GRP170, reducing overall protein translation, and expanding ER volume(41). Beyond these responses, downstream, UPR effectors augment protein folding and the destruction of misfolded proteins via ERAD, thereby reducing the protein load in the ER (29, 42). If ER proteostasis is not restored or stress persists, however, the UPR instead induces pro-apoptotic transcription factors, such as CHOP, which ultimately initiate a cell death cascade(42, 43). Indeed, we showed that induced depletion of GRP170 in the nephron of the GRP170^NT-/-^ mouse was associated with UPR activation and cell death. We therefore proposed that the UPR triggers the AKI-like phenotypes noted above (20). Nevertheless, whether renal dysfunction and kidney injury in our GRP170^NT-/-^ mouse model was a direct consequence of UPR activation was unclear.

To potentially uncouple renal damage from the UPR upon GRP170 deletion *in vivo*, and more generally to better define the role of ER chaperone function in AKI, we subjected GRP170^NT-/-^ mice and littermate controls to a high sodium (8%) diet (HS) after dox-inducible KO of the *HYOU1* loci. The basis of this approach is that extra dietary sodium should compensate for the impaired water and electrolyte homeostasis we previously observed in GRP170-deficient animals (20). Here, we show that repleting lost electrolytes improves intravascular volume and ameliorates the AKI phenotype, but ultimately fails to prevent renal damage and pro-apoptotic UPR activation. Together, these data support the role of the UPR in epithelial cell homeostasis in the nephron, highlight the critical function of the GRP170 chaperone in renal function, and support the use of high sodium diets—in combination with drugs that temper UPR activation and restore ER function—in AKI(44, 45).

## METHODS

### Animal maintenance

Inducible renal-tubule-epithelial-specific GRP170^flox/flox^/Pax8-rTA/LC1 knockout mice (GRP170^NT-/-^) were generated in a C57Bl/6 background as previously described(20, 46). Age-matched GRP170^NT-/-^ littermates (GRP170^flox/flox^/WT/LC1 or GRP170^flox/flox^/Pax8-rTA/WT) were used as controls. All experiments conformed to NIH Guide for the Care and Use of Laboratory Animals and were approved by the University of Pittsburgh IACUC. Male and female mice (average age 22 weeks) were used in each experiment. Mice were housed in a temperature-controlled room with a 12 hour light-dark cycle and allowed free access to deionized water. To induce gene deletion, doxycycline (0.2% in sucrose) was added to the drinking water for 10 days as previously described(20). Gene deletion was confirmed by qPCR of samples purified from whole kidneys.

To assess the effect of excess dietary sodium on renal physiology, mice were provided either standard chow (Prolab Isopro RMH 3000) or high sodium (HS) chow (Teklad diet, Indianapolis, IN) from the start of doxycycline administration until the animals were sacrificed. Standard and HS chow contains 0.3% and 8% sodium, respectively. Animals were weighed a minimum of every 3 days.

### Serum chemistry

Whole blood was aspirated from the right ventricle of anesthetized mice. Blood chemistry (Na^+^, K^+^, Cl^-^, BUN, creatinine, glucose, hemoglobin, and hematocrit) was assessed using an iStat (Abbott Point of Care). The upper limit of quantification for BUN is 140 mg/dl and the lower limit of detection for creatinine is 0.2 mg/dl. For statistical validation, values above or below these thresholds were considered 140 mg/dl or 0.2 mg/dl, respectively. Plasma aldosterone concentration was assessed using the Enzo Life Sciences Aldosterone ELISA kit (ADI-900-173) per the manufacturer’s instructions.

### Metabolic cage measurements

To assess water intake, urine composition and urine volume, mice were individually housed in metabolic cages (Tecniplast) and fed gel-based diet prepared from either standard or HS chow as described above. After allowing at least 24 hours to acclimate to the cage and food, 24-hour water consumption and urine volume were recorded for two consecutive days. Urine osmolality was measured using an Osmette Micro-Osmotte Osmometer (Precision Systems). Urine Na^+^, K^+^, and glucose were measured using flame photometry.

### RNA extraction and quantitative PCR (qPCR)

Total RNA was extracted from 25 mg of whole kidney tissue by manual homogenization using a 25-gauge needle in RLT buffer as described by the manufacturer (Qiagen RNeasy Mini Kit). The purity of RNA was evaluated using a spectrophotometer (Thermo Fisher Scientific, NanoDrop One Microvolume UV-Vis Spectrophotometer) and quality confirmed with agarose gel electrophoresis. cDNA was amplified from 1 μg of total RNA according to the manufacturer’s instructions (QuantaBio qScript Supermix.) cDNA concentration was measured by spectrophotometry, and samples diluted to a concentration of 15 ng/μL. Using 80 ng of cDNA per replicate, quantitative PCR (qPCR) was carried out using the QuantStudio III system (Thermo Fisher Scientific) as previously described. Relative amplification was calculated by averaging results from a minimum of three technical replicates and three to five biological replicates. Gene expression levels were normalized to β-actin, and fold changes in gene expression were calculated using the 2^(-ΔΔCt) method.

### Kidney histology and TUNEL Staining

Kidneys were harvested and preserved in formalin (for IHC) or 4% PFA (for immunofluorescence). Fixed kidneys were embedded in paraffin and sectioned at a thickness of 4 µm for either periodic Acid-Schiff (PAS) or terminal deoxynucleotidyl transferase biotin-dUTP nick-ended labeling (TUNEL) staining using standard histological procedures. To assess renal histology, whole kidney sections, including the cortex and medulla, were scanned with a 20x objective on a EVOS FL Auto Cell Imaging System (Thermo Fisher Scientific) microscope. Apoptotic nuclei were identified by TUNEL staining using the ApoTag Fluorescein In situ Apoptosis Detection Kit S7110 (MilliporeSigma) according to the manufacturer’s instructions. Tissues were observed under a microscope Leica TCS SP8 Confocal Microscope (Leica Microsystems).

### Statistics

Results are presented as mean +/- standard deviation (SD). Graphpad Prism software (version 9) was used for statistical analysis. When comparing two groups, a Students’ t-test was performed. For comparison of three or more experimental groups, ANOVA with *post-hoc* Tukey multiple comparison tests or Brown-Forsythe ANOVA with *post-hoc* Benjamini multiple comparison tests (mRNA expression) were used to assess significance. In all cases, a threshold of p <0.05 was considered statistically significant.

## RESULTS

### A high sodium diet normalizes serum electrolytes and improves intravascular volume in GRP170 KO mice

Our first objective was to better define the role of GRP170 in electrolyte homeostasis. Therefore, we deleted GRP170 from the nephron by administering dox in drinking water to GRP170^NT-/-^ and control mice for 10 days, as described previously (20). Simultaneously, mice were fed either a HS diet (8% sodium), or a standard diet (0.3% sodium) for 21 days and then sacrificed (see Fig. 1A). We previously showed that levels of GRP170 message and protein in the GRP170^NT-/-^ mice fell to ∼25% of the control, and the animals had lost ∼20% of their initial body weight by day 21 post-dox treatment (20). Thus, we predict that this would maximize our ability to observe rescue by a HS diet and allow for a direct comparison to our extensive previous work(20). Because of the extreme volume and hyponatremia previously described for GRP170^NT-/-^ mice, we chose to administer the 8% sodium diet rather than a more moderate sodium diet. In addition, high sodium diets are used clinically for patients exhibiting hyponatremia or low blood pressure(47). Unless otherwise specified, both male and female animals were used in each of the ensuing experiments (see below).

**Figure 1:**
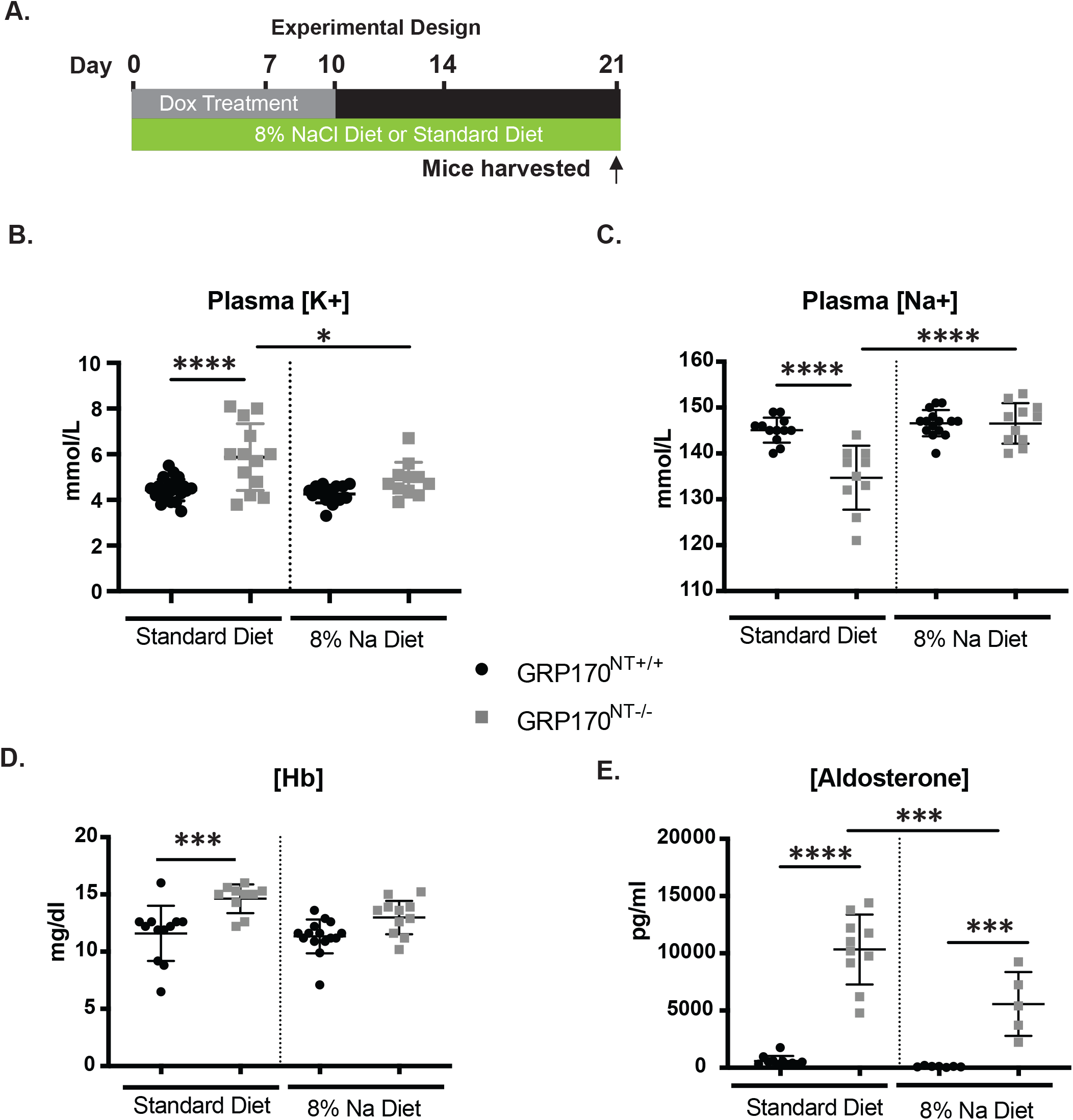
High sodium diet partially restores intravascular volume depletion and normalizes plasma sodium and potassium concentrations in mice lacking GRP170 expression in the nephron tubular epithelium. (A) Mice were treated with doxycycline to induce GRP170 deletion and simultaneously fed a standard chow or 8% sodium diet until sacrifice after 21 days. Blood chemistry data is presented here (B and C) plasma electrolytes, (D) hemoglobin, and (E) aldosterone. Data represent the means +/- SD; n= 4-13. *P<0.05, **P < 0.01, ***P < 0.001, ****P < 0.0001. Statistical significance determined by 1-way ANOVA followed by Tukey’s multiple-comparison test.

First, we observed that GRP170^NT-/-^ mice on a HS diet maintained serum sodium and potassium concentrations comparable to control mice on either the HS or standard diet (Fig. 1B and C). In contrast and consistent with our prior data (20), the GRP170^NT-/-^ mice on the standard diet developed hypokalemia and hyponatremia. Moreover, the HS diet normalized hemoglobin levels of the GRP170^NT-/-^ mice (Fig. 1D). These combined results suggest that additional sodium helps preserve intravascular volume and overcomes the natriuretic effect caused by GRP170 depletion. However, GRP170^NT-/-^ mice on the HS diet still exhibit significant weight loss (Supplemental Fig. S1A). The observed weight loss is consistent with persistent, mild volume depletion and excess urinary water excretion (see below). Formally, weight loss could also reflect a catabolic state induced by GRP170 deletion (see Discussion).

Secretion of serum aldosterone is stimulated primarily by hyperkalemia and to a lesser extent by volume depletion (16). Based on the normalization of plasma sodium, potassium, and hemoglobin levels in GRP170^NT-/-^ mice on a HS diet, we predicted that aldosterone levels would be lower in GRP170^NT-/-^ mice fed a HS rather than the standard diet. Consistent with this hypothesis, extra dietary sodium reduced aldosterone levels in GRP170^NT-/-^ mice, yet aldosterone remained significantly higher in GRP170^NT-/-^ mice than in control animals fed either diet (Fig. 1E). The most likely explanation for this result is that volume depletion persists.

### A high sodium diet fails to correct urinary electrolyte and water handling

We next hypothesized that increased plasma aldosterone (Fig. 1F) represents an attempt to maintain potassium homeostasis and/or intravascular volume, potentially due to impaired ion or water channel expression. Yet, renal electrolyte homeostasis remains compromised in GRP170^NT-/-^ mice even when given a HS diet. In fact, this is consistent with ongoing weight loss in GRP170^NT-/-^ mice on the HS diet (Fig. S1). Hence, we next assessed renal water and electrolyte handling using metabolic cages to measure urine volume, osmolality, and composition, and water intake of individual animals.

Typically, after a brief period of re-equilibration, during which extra dietary sodium and, thus, water is reabsorbed, daily sodium intake equals excretion. Therefore, as expected based on the pronounced weight loss in the GRP170-depleted animals, control and GRP170^NT-/-^ animals on a HS diet produced more urine (Fig. 2A), consumed more water (Fig. 2B), and excreted more sodium (Fig. 2C) than littermates of the same genotype fed a standard diet. However, GRP170^NT-/-^ mice fed HS chow still had significantly higher 24-hour urine output, as well as lower urine osmolality, compared to control littermates on the HS diet (Fig. 2A, D). The GRP170^NT-/-^ animals also excreted less sodium and potassium than controls (Figures 2C, E). Because the most efficient means of preserving intravascular volume is to retain sodium and water, GRP170^NT-/-^ mice on a HS diet are also expected to produce a relatively more concentrated urine than those on a standard diet, provided functional ion channels and transporters are present at the epithelial cell apical membrane (17). The fixed and relatively low urine osmolality of GR170 deficient animals (Fig. 2D) indicates, therefore, that a HS diet does not reverse the severe urine concentrating defect. This may be a consequence of damaged PT cells, impaired sodium reabsorption in the loop of Henle and distal collecting duct, and/or diminished water transport in the collecting duct (17).

**Figure 2:**
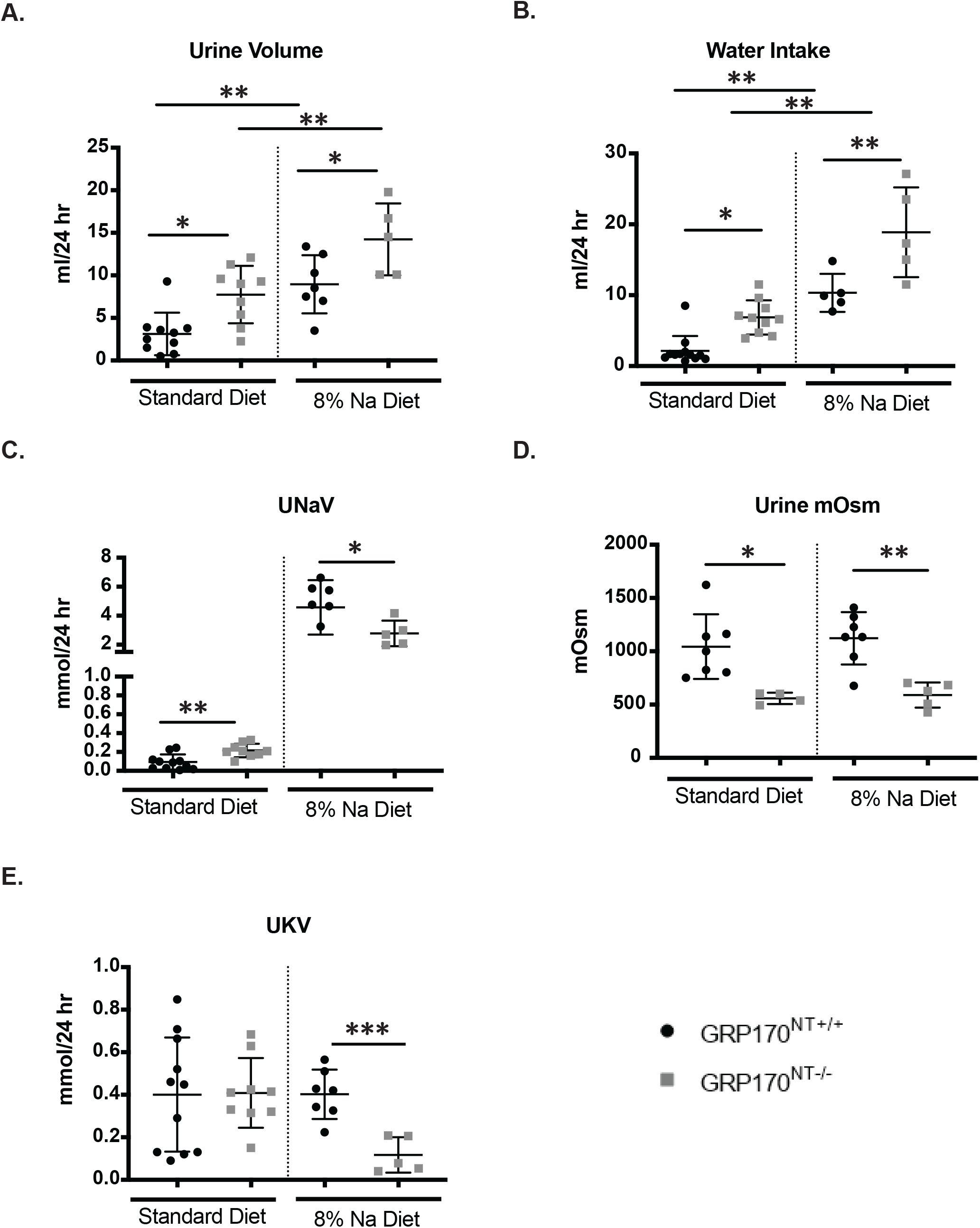
High sodium diet fails to rescue the urine concentrating defect and naturesis of GRP170^NT-/-^ mice. Mice were kept individually in metabolic cages to allow collection of 24 hour urine samples. Urine samples were collected and water consumption measured on the day prior to sacrifice. (A)Urine volume, (B) water intake and (C) urine osmolality; (E and F) 24 hour sodium and potassium excretion; Data represent the means +/- SD; n= 4-10. *P<0.05, **P < 0.01, ***P < 0.001, ****P < 0.0001. Pairwise comparisons were made for (C).

### The high sodium diet also ameliorates kidney injury in the GRP170^NT-/-^ mice

Our results suggest that extra dietary sodium partially restores intravascular volume in the dox-treated GRP170^NT-/-^ animals. Because impaired renal perfusion resulting from volume depletion can cause AKI, we predicted that GRP170-depleted animals fed a HS diet will present with a milder kidney injury phenotype than GRP170^NT-/-^ mice on a standard diet as well as the WT littermates, regardless of their diet(48). To test this hypothesis, we first measured the levels of two established AKI biomarkers, BUN and creatinine(48).

Consistent with reduced injury, the cohort of GRP170^NT-/-^ mice on a HS diet had significantly lower serum BUN and creatinine than those on a standard diet (Fig. 3A, B), and BUN concentration in GRP170^NT-/-^ mice was no longer significantly elevated relative to control animals fed either diet. Furthermore, when we assessed albuminuria, a marker of PT dysfunction (7, 49), the amount of albumin in the urine from GRP170^NT-/-^ mice on a HS diet, while still elevated, were somewhat lower than those on a standard diet (compare Fig. 3C to published data(20). As expected, minimal levels of albumin were found in urine of control littermates on the standard or HS diet. Next, to detect more subtle and specific evidence of epithelial cell injury in the nephron, we quantified message levels corresponding to the tubular injury marker, NGAL (50, 51), from whole kidney specimens. As shown in Fig. 3D, NGAL transcripts were elevated in GRP170^NT-/-^ animals versus littermate controls, regardless of diet and—although not statistically significant—were further elevated in animals fed a HS diet. This latter result which may reflect the inherent nephrotoxicity of excess dietary sodium (see below) (52). In aggregate, these findings suggest that a HS diet modestly improves but does not prevent select AKI-like features in GRP170^NT-/-^ mice.

**Figure 3:**
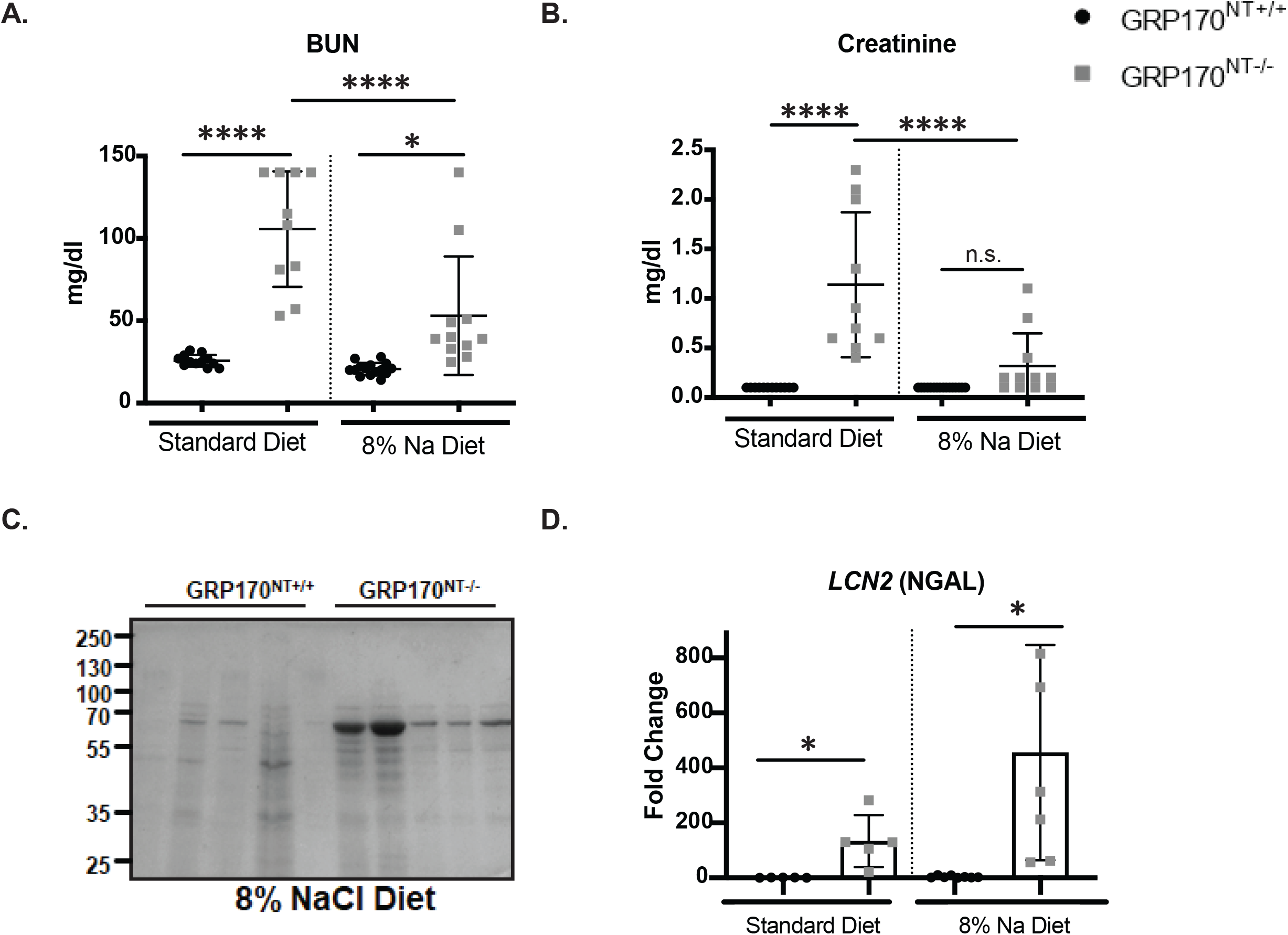
A high sodium diet ameliorates but does not prevent kidney injury due in GRP170^NT-/-^ mice. (A and B) Serum BUN and creatinine concentration. (C) Equal volumes of urine were subjected to SDS-PAGE and stained with Coomassie blue to detect urinary albumin excretion. (D) *LCN2* (NGAL) relative mRNA expression. Data represent the means +/- SD; n= 4-10. *P<0.05, **P < 0.01, ***P < 0.001, ****P < 0.0001.

### Histopathologic evidence of AKI in GRP170^NT-/-^ mice is unaltered by a high sodium diet

In individuals with AKI or in established AKI mouse models, evidence of renal damage can be visualized with Periodic Acid-Schiff staining (PAS). Therefore, we next examined histologically the extent to which a HS diet rescues kidney injury in the GRP170^NT-/-^ mice. PAS-staining of kidney sections revealed that a HS diet had minimal effects on the degree of tubular dilation, cell volume, brush border loss, intraluminal cell sloughing, and cast formation in GRP170-depleted mice compared to GRP170^NT-/-^ mice on a control diet (Fig. 4). In contrast to the kidney injury markers examined in Fig. 3, PAS staining of kidney sections from WT littermates fed either the control or HS diet were indistinguishable. Interestingly, the histopathologic features in the GRP170^NT-/-^ mice remained more prominent in the outer cortex than in the medulla. Since PTs predominate in the cortex, whereas other nephron segments are found primarily in the medulla (4), the injury pattern illustrates the vital role of GRP170—and ER proteostasis more generally—in PT epithelial cell health. These results suggest that the PT is the site of initial injury. This conclusion is especially striking since the cortex is better perfused and, consequently, better positioned to benefit from volume expansion than the medulla (53).

**Figure 4:**
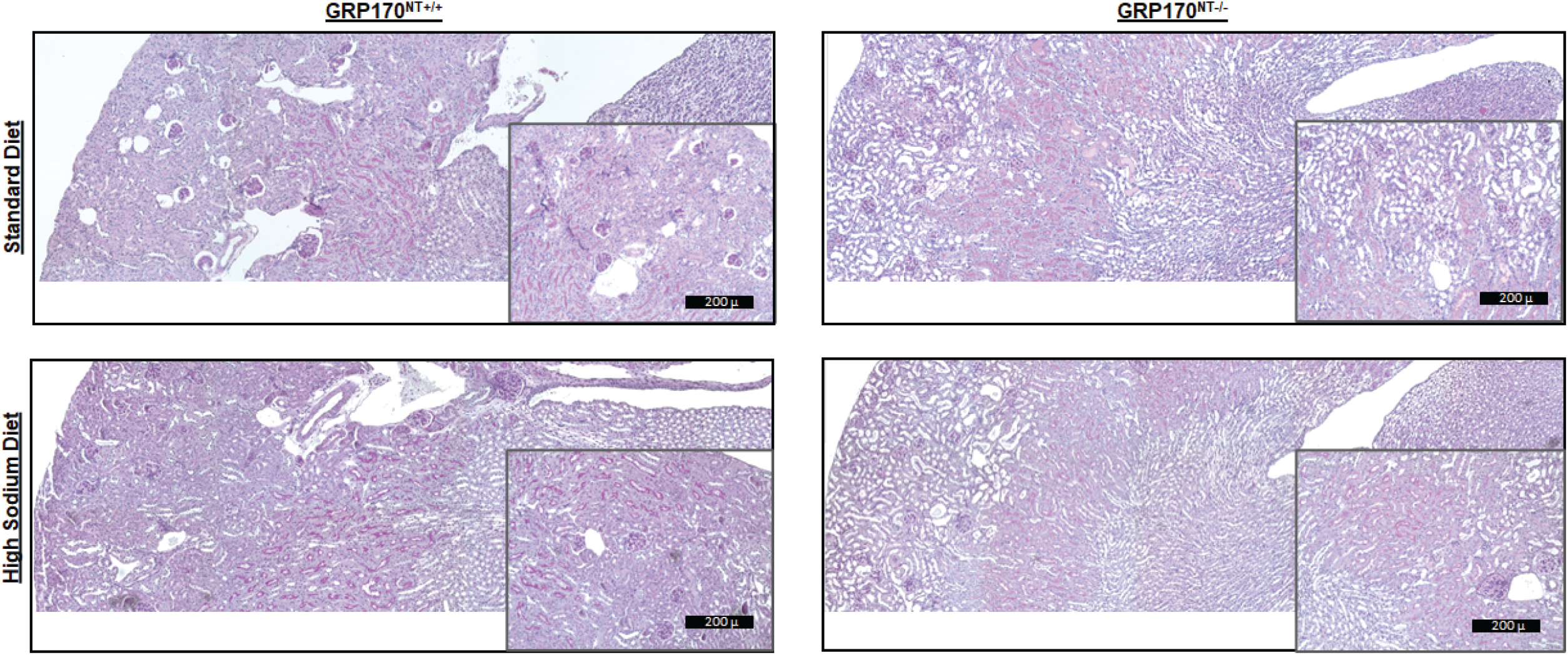
High sodium diet does not prevent kidney injury caused by loss of GRP170 expression in the nephron epithelium. Representative PAS stained kidney sections from female GRP170^NT-/-^ and GRP170^NT-/-^ mice fed either a standard or an 8% sodium diet are shown. GRP170^NT-/-^ mice exhibit histological findings characteristic of kidney injury: tubular epithelial thinning and dilation, epithelial cell sloughing, and granualar casts. Scale bar: 200 microns.

### High sodium diet fails to decrease the UPR in GRP170^NT-/-^ mice

We previously demonstrated that progressive loss of GRP170 is accompanied by induction of the UPR in GRP170^NT-/-^ mice (20). UPR activation and GRP170 expression have also been linked to numerous disease states, including ischemic injury, diabetes, diabetic nephropathy, cancer, and neurodegenerative disease (27, 32, 54–58). We also recently showed that the progressive loss of GRP170 is accompanied by UPR activation and induction of apoptotic markers in mouse embryo fibroblasts (38). Therefore, we asked if a HS diet tempers UPR activation as a result of the downstream consequence of the amelioration of select kidney injury markers in GRP170^NT-/-^ mice (see above). To address this question, we compared steady-state mRNA levels associated with UPR-associated genes induced by each of the three UPR branches: sXbp1, which is downstream of IRE1 activation, BiP, which is downstream of ATF6 activation, and CHOP, which is downstream of PERK activation(59). The levels of each were measured by qPCR in isolated kidneys from the GRP170^NT-/-^ mice fed either a HS or standard diet. Consistent with loss of GRP170 activating a global UPR, the levels of all three markers rose significantly in the GRP170^NT-/-^ mice fed a standard diet compared to their WT littermates (Fig. 5A-C). We also found that the HS diet was unable to blunt the elevated UPR and, in fact, sXbp1, and CHOP levels were even higher in animals fed a HS diet, suggesting secondary effects on ER function in cells exposed to high salt, as suggested by previous studies (60–63) (also see Discussion).

**Figure 5:**
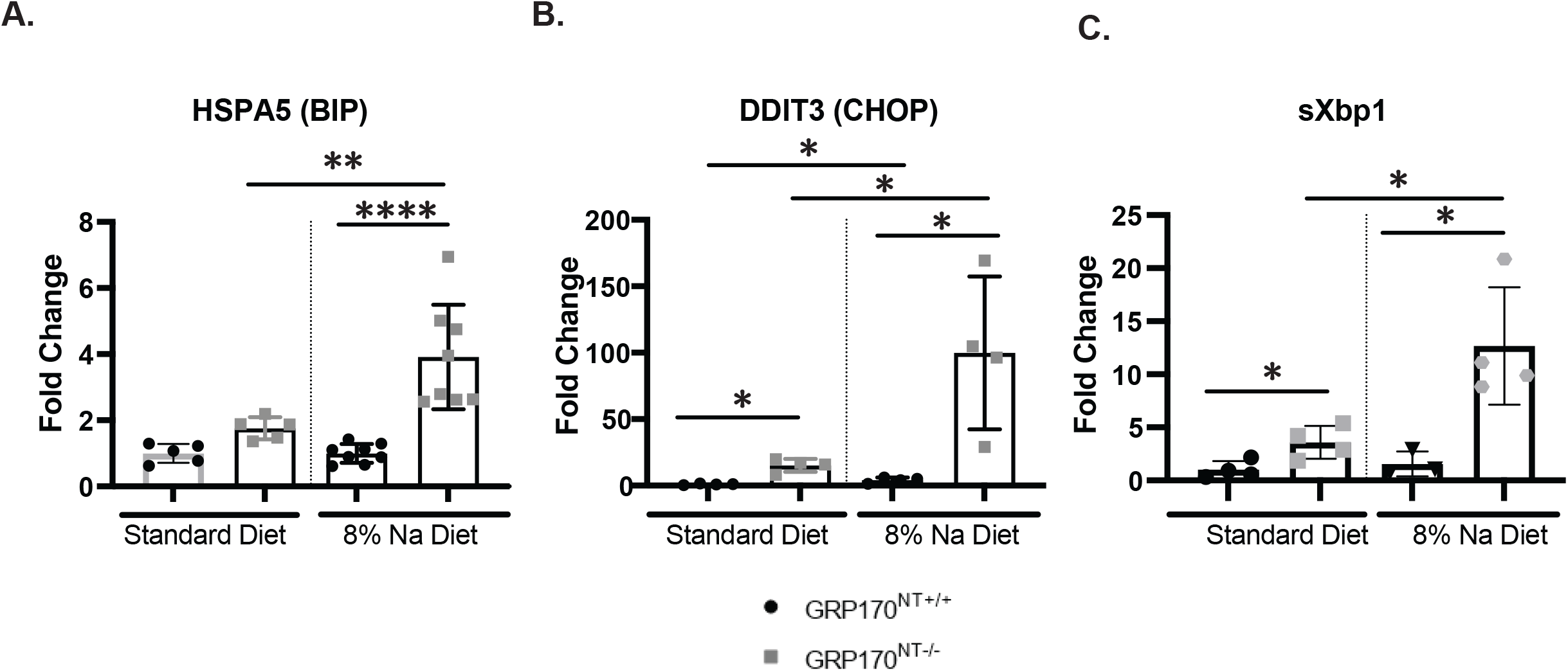
The unfolded protein response (UPR) is induced to a similar degree in GRP170^NT+/-^ animals fed a standard and high sodium diet. (A, B, C) mRNA transcript expression of UPR effectors determined by qPCR from whole kidneys from male mice. Data represent the means +/- SD; n= 4-10. *P<0.05, **P < 0.01, ***P < 0.001, ****P < 0.0001.

It is also important to note that BiP expression is temporally related to ER stress, with transient induction followed by gradual resolution (64). We previously noted that BiP mRNA expression peaked at day 14 post dox and then recovered by day 21 in dox-fed GRP170^NT-/-^ mice (20). Hence, elevated BiP mRNA observed in the HS diet cohort, in combination with milder renal injury (see above), is consistent with the possibility that that the HS diet delays activation of the IRE1 arm of the UPR. Unfortunately, it is impossible to test this hypothesis since the GRP170^NT-/-^ mice have lost ∼20% of their bodyweight by day 21 and must be sacrificed.

Because chronic UPR activation induces apoptotic cell death, we next asked if the HS diet ameliorated apoptosis in GRP170 deficient animals(59). As shown in Fig. 6, however, renal cells examined from GRP170^NT-/-^ mice had a comparable number of TUNEL positive nuclei regardless of diet, while those from GRP170^+/+^ mice had few TUNEL positive cells. The presence of TUNEL positive cells and elevated expression of the pro-apoptotic transcription factor, CHOP, in GRP170^NT-/-^ animals suggest that the HS diet is unable to maintain cell viability. Overall, these data suggest that compromised renal electrolyte handling drives a portion, but not all, of the AKI-like phenotypes of the GRP170-deficient mice. Our results also suggest UPR-mediated tissue damage is also a likely contributor to kidney injury.

**Figure 6:**
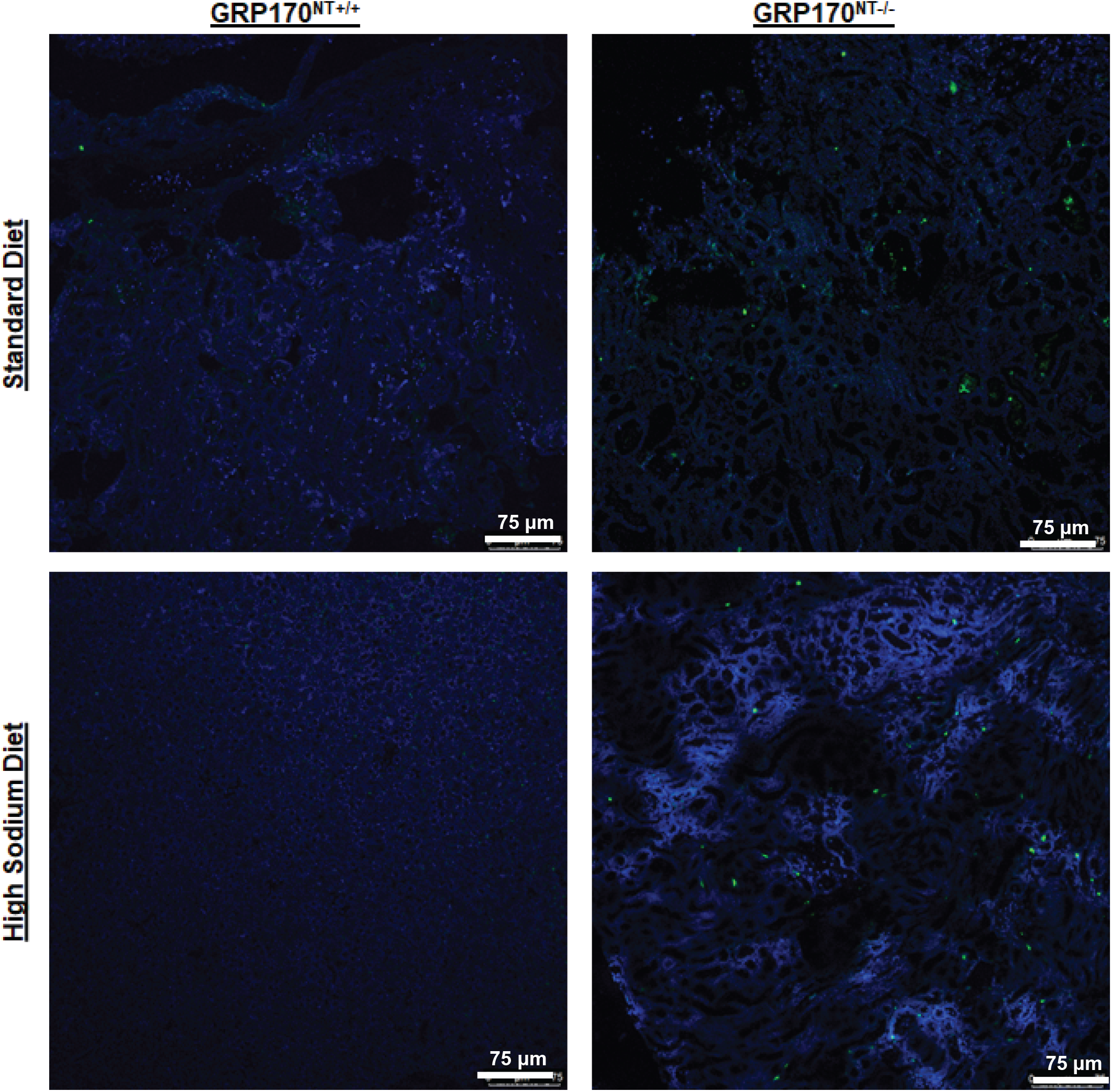
High sodium diet does not ameliorate renal tubular apoptosis caused by loss of GRP170. Representative staining of kidney sections from GRP170^NT-/-^ and GRP170^NT-/-^ male mice fed eith a standard or an 8% sodium diet are shown. Scale bar: 75 microns.

### Male GRP170^NT-/-^ mice are more refractory to high sodium restoration of select phenotypes

Sex specific differences in renal physiology are well-documented. For example, males of both humans and rodents are more sensitive to kidney injury than females (65–68). We previously determined that male GRP170^NT-/-^ mice were also more sensitive to the loss of GRP170 than female mice (20). To assess whether sex-dependent effects also correlated with the extent of rescue when GRP170^NT-/-^ mice were maintained on a HS diet, we reanalyzed the data for several phenotypes by sex. In brief, we found that the HS diet normalized both plasma sodium and potassium (Fig. 7A-B) and led to volume expansion, as indicated by hemoglobin levels (Fig. 7C) in both sexes. In contrast, the HS diet failed to rescue GRP170 depletion-associated weight loss in either sex (Fig. S1). As we previously found, male GRP170^NT-/-^ mice were again more sensitive to injury, as reflected in the BUN and creatinine levels (20) (Fig. 7D-E). Strikingly, however, BUN and creatinine levels were normalized in female mice maintained on a HS diet, whereas the levels of these kidney injury markers, while lower, remained elevated in male mice. Renal histology revealed that male mice additionally exhibited more extensive tubular injury than females, though the effect of dietary sodium supplementation was negligible in both cases (Fig. 4 and Fig. S2). Therefore, we conclude that the HS diet more completely rescues the GRP170 KO in female than male mice.

**Figure 7:**
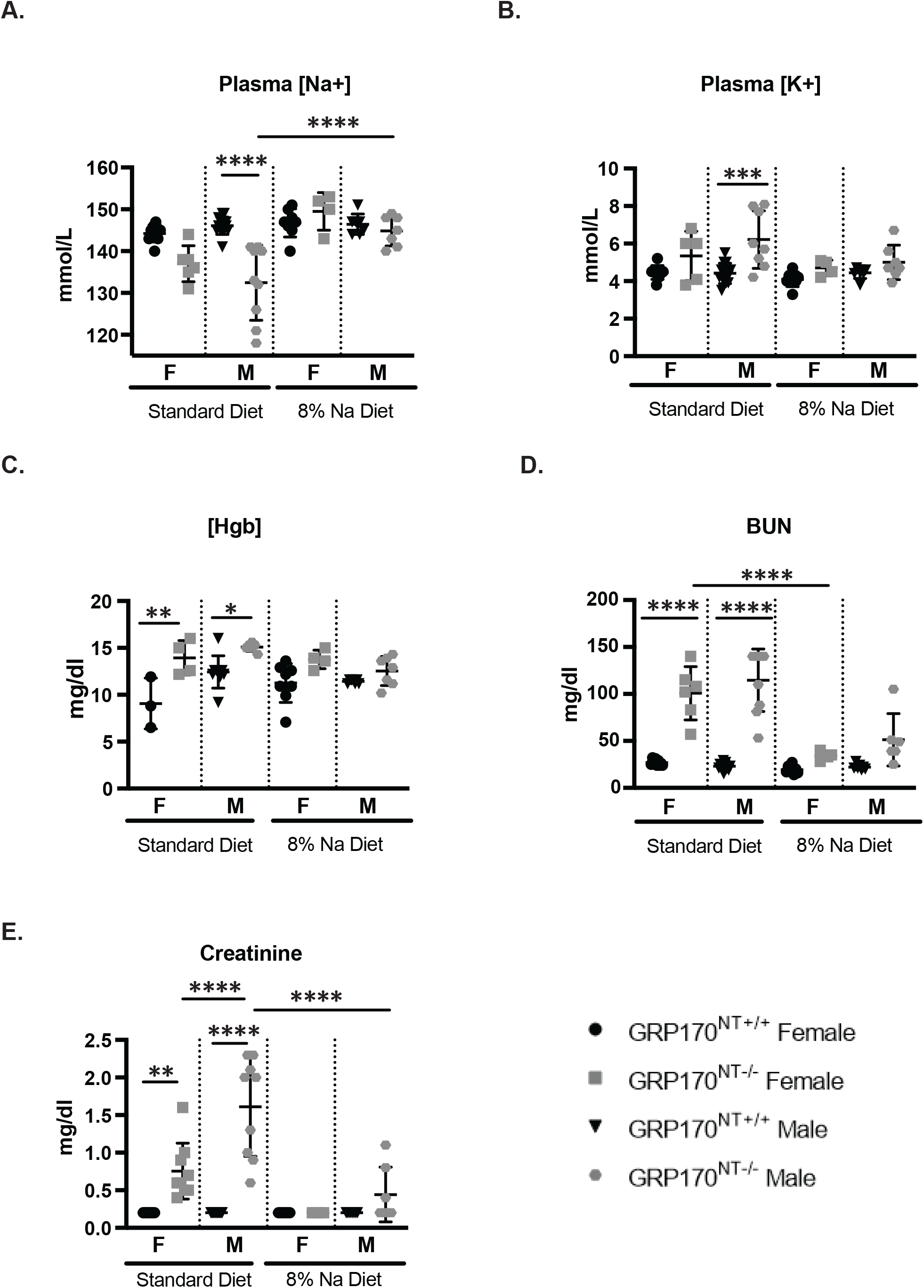
High sodium diet restores resuces female GRP170^NT-/-^ mice more completely than male GRP170^NT-/-^ mice. (A and B) Plasma electrolytes, (C) hemaglobin, (D) BUN and (E) creatinine. Data represent the means +/- SD; n= 4-13. **P < 0.01, ***P < 0.001, ****P < 0.0001. Statistical significance determined by 1-way ANOVA followed by Tukey’s multiple-comparison test (3 or more data sets).

## DISCUSSION

Molecular chaperones in the ER are vital for protein folding, post-transcriptional modifications, the removal of defective proteins, and assembly of secreted and transmembrane proteins, and defects in these processes have been linked to many diseases (18, 19, 69, 70). Our lab previously demonstrated that one such chaperone, GRP170, is required for the biogenesis of ENaC (Buck 2013, Buck 2017, Buck 2023), and that GRP170 deletion in the nephron activates the UPR and leads to an AKI-like phenotype (20). However, the mechanism by which GRP170 contributes to nephron epithelial integrity is unknown. Because a high sodium diet augments intravascular volume and substantially ameliorates the renal phenotype of ENaC and mineralocorticoid receptor KO mice, we hypothesized that supplemental sodium might also compensate for GRP170 deletion by overcoming the inappropriate naturesis and water loss that accompanies GRP170 deficiency (20, 34, 35, 37, 71). Here, we report that sodium supplementation restored intravascular volume and corrected hyponatremia and hyperkalemia and ameliorated AKI, but failed to significantly blunt UPR activation in GRP170-deficient mice. These results strongly suggest that UPR activation also contributes to AKI in the GRP170 depletion model.

Similar to sodium supplementation in GRP170^NT-/-^ mice, sodium supplementation in α, β and γENaC and mineralocorticoid receptor deletion mice rescued many of the phenotypes associated with these model systems (34–36, 71). In particular, we report that HS intervention improved intravascular volume, normalized serum electrolytes, and somewhat reduced plasma aldosterone levels. However, sodium supplementation was unable to prevent weight loss in GRP170^NT-/-^ mice or reverse the urine concentrating defect and naturesis, as it did in the mineralocorticoid and αENaC KO models (35, 71). Since intravascular volume appears restored in GRP170^NT-/-^ mice, ongoing weight loss may reflect—at least in part—a refractory catabolic state linked to GRP170 deficiency. Notably, previous studies reported some Pax8 expression in thyroid tissue(72, 72), which in principle may contribute to weight loss in GRP170^NT-/-^ animals (Fig. S1). Nevertheless, GRP170^NT-/-^ mice are also distinguished by their AKI-like phenotype. Though saline loading partially ameliorated kidney injury, GRP170 deficient mice still had refractory azotemia and elevated NGAL expression. PAS staining also revealed pathologic hallmarks of AKI.

The persistence of urinary abnormalities and an AKI-like phenotype in GRP170^NT-/-^ mice points to a crucial, generalized role for GRP170 in renal physiology beyond its effects on distal tubular sodium handling. This conclusion is supported by persistent albuminuria, an indication of PT dysfunction, in these animals(48).

Histologic evidence of persistent severe injury in the renal cortex, and relative sparing of the medulla despite volume expansion, further suggests that injury occurs initially in the PT rather than in the distal nephron. Thus, GRP170 deficiency directly affects PT homeostasis. Consequently, we propose that proximal tubulopathy is likely the dominant phenotype of GRP170 depleted mice because the PT is highly dependent on aerobic metabolism and poorly perfused, even under ideal circumstances (14). Hence, the PT is especially vulnerable to oxygen debt or other cell stressors, such as the UPR and ER stress, which can also compromise mitochondrial function and impair ATP generation (53, 73–75).

By enhancing renal perfusion, supplemental sodium may have delayed kidney injury without addressing a secondary source of injury: impaired ER proteostasis and UPR activation. This conclusion is consistent with the observation that expression of a pro-apoptotic transcription factors, CHOP, was induced in GRP170-depleted animals given a high-salt diet. Activation of the UPR-transducer, PERK, leads to CHOP induction(76). While PERK signaling is transiently cytoprotective, sustained PERK instead initiates CHOP-mediated apoptosis (42, 64). In addition, TUNEL positive, apoptotic, cells persist in GRP170-deficient animals despite a HS diet. Paradoxically, and as documented in our prior work (20), markers of the adaptive UPR, including BiP and Xbps1, were also upregulated. This may reflect the myriad functions of GRP170 as both a chaperone and integral element of the UPR (also see below) (77).

Our analysis of UPR activation in animals fed a high-sodium diet is complicated by the effects of sodium on cell stress. Excess sodium consumption leads to oxidative injury and hypoxia, and can predispose animals to AKI (78, 79). Moreover, hyperosmotic environments alter membrane protein trafficking and favor protein aggregation, thereby causing ER stress and activating the UPR (60–63). This is especially relevant since the nephron is exposed to wide fluctuations in osmolality. For example, in a collecting duct principal cell model, NaCl administration preferentially upregulated BiP and ATF6 expression and PERK phosphorylation, but not XBP1 splicing (63), a pattern which favors cell death rather than resilience. In addition, rats administered 1.5% NaCl in drinking water for eight weeks had elevated urinary Kim1 levels, and ATF4 and CHOP expression.

Transcription of Bcl-2, an inhibitor of apoptosis, was also reduced. Interestingly, though, there was evidence of mild tubular dilation, inflammation, and fibrosis in kidneys obtained from these animals, their serum BUN and creatinine concentrations were indistinguishable from mice provided distilled water (80). In our current study, we similarly noted higher CHOP levels in control animals fed the HS compared to standard diet. The combined effect of GRP170 deficiency and sodium stress may explain why GRP170^NT-/-^ mice fed a HS diet exhibited elevated UPR induction and NGAL/Kim1 expression than those fed a standard diet. While the confounding effects of sodium limit conclusions about the pattern of UPR activation, the dramatic UPR induction seen in GRP170^NT-/-^ mice nonetheless highlights the chaperone’s profound importance in the UPR.

As with diet, sex influences renal physiology and, quite likely, the UPR as well. In mice and humans, females are less susceptible to kidney injury (81–84). Each sex also regulates sodium reabsorption differently. For example, female mice excrete a saline load more rapidly than males because of reduced proximal tubule reabsorption (85, 86). Yet, the basal levels of NCC and cleaved α and γENaC expression are as much as 50% higher than males, potentially facilitating more effective distal sodium reabsorption and potassium excretion (85). Finally, males are more sensitive to ER stress-related kidney injury and exhibit greater induction of BiP, CHOP, and XBP1s compared to females (67). Our results are consistent with these observations. In the case of GRP170^NT-/-^ mice, female mice likely have a milder phenotype than males because of a combination of reduced vulnerability to AKI, more baseline ENaC expression, and—probably most importantly—a less robust UPR. Critically, the relatively modest UPR and AKI phenotypes of females lends credence to the potential for a genetic or epigenetic UPR suppressor which ameliorates kidney injury.

In conclusion, this study further establishes our inducible murine model as a powerful tool with which to probe the relationship between the UPR, molecular chaperone function, and AKI, and to clarify which downstream UPR pathways and factors promote and/or kidney injury. Based on the partial rescue of several phenotypes associated with the emerging links between these phenomena, it will be critical in future work to determine whether UPR-directed pharmaceuticals can prevent or ameliorate AKI, and whether their effects are magnified when combined with a HS diet.

## GRANTS

This work was supported by NIH grant R01 DK117126 (T.M.B.), NIH grant R35 GM131732 (J.L.B.), an NIH training grant T32 K091202 (A.P.) and NIH grant K12 (NICHD). Institutional support from NIH grant U54 DK137329 is also acknowledged.

## REFERENCES

1. Susantitaphong P, Cruz DN, Cerda J, Abulfaraj M, Alqahtani F, Koulouridis I, Jaber BL. World Incidence of AKI: A Meta-Analysis. Clin J Am Soc Nephrol 8: 1482–1493, 2013. doi: 10.2215/CJN.00710113.

2. Coca SG, Yusuf B, Shlipak MG, Garg AX, Parikh CR. Long-term Risk of Mortality and Other Adverse Outcomes After Acute Kidney Injury: A Systematic Review and Meta-analysis. American Journal of Kidney Diseases 53: 961–973, 2009. doi: 10.1053/j.ajkd.2008.11.034.

3. Basile DP, Anderson MD, Sutton TA. Pathophysiology of Acute Kidney Injury. Compr Physiol 2: 1303–1353, 2012. doi: 10.1002/cphy.c110041.

4. Jacobson HR. Functional segmentation of the mammalian nephron. Am J Physiol 241: F203–218, 1981. doi: 10.1152/ajprenal.1981.241.3.F203.

5. Sandoval RM, Wagner MC, Patel M, Campos-Bilderback SB, Rhodes GJ, Wang E, Wean SE, Clendenon SS, Molitoris BA. Multiple Factors Influence Glomerular Albumin Permeability in Rats. J Am Soc Nephrol 23: 447–457, 2012. doi: 10.1681/ASN.2011070666.

6. Tenten V, Menzel S, Kunter U, Sicking E-M, van Roeyen CRC, Sanden SK, Kaldenbach M, Boor P, Fuss A, Uhlig S, Lanzmich R, Willemsen B, Dijkman H, Grepl M, Wild K, Kriz W, Smeets B, Floege J, Moeller MJ. Albumin Is Recycled from the Primary Urine by Tubular Transcytosis. J Am Soc Nephrol 24: 1966–1980, 2013. doi: 10.1681/ASN.2013010018.

7. Curthoys NP, Moe OW. Proximal Tubule Function and Response to Acidosis. Clin J Am Soc Nephrol 9: 1627–1638, 2014. doi: 10.2215/CJN.10391012.

8. Moe OW, Ujiie K, Star RA, Miller RT, Widell J, Alpern RJ, Henrich WL. Renin expression in renal proximal tubule. J Clin Invest 91: 774–779, 1993. doi: 10.1172/JCI116296.

9. Legouis D, Faivre A, Cippà PE, de Seigneux S. Renal gluconeogenesis: an underestimated role of the kidney in systemic glucose metabolism. Nephrology Dialysis Transplantation 37: 1417–1425, 2022. doi: 10.1093/ndt/gfaa302.

10. Chesney RW. Interactions of vitamin D and the proximal tubule. Pediatr Nephrol 31: 7–14, 2016. doi: 10.1007/s00467-015-3050-5.

11. Kurtz I. Renal Tubular Acidosis: H+/Base and Ammonia Transport Abnormalities and Clinical Syndromes. Advances in Chronic Kidney Disease 25: 334–350, 2018. doi: 10.1053/j.ackd.2018.05.005.

12. Scholz H, Boivin FJ, Schmidt-Ott KM, Bachmann S, Eckardt K-U, Scholl UI, Persson PB. Kidney physiology and susceptibility to acute kidney injury: implications for renoprotection. Nat Rev Nephrol 17: 335–349, 2021. doi: 10.1038/s41581-021-00394-7.

13. Ho KM, Morgan DJR. The Proximal Tubule as the Pathogenic and Therapeutic Target in Acute Kidney Injury. NEF 146: 494–502, 2022. doi: 10.1159/000522341.

14. Cargill K, Sims-Lucas S. Metabolic requirements of the nephron. Pediatr Nephrol 35: 1–8, 2020. doi: 10.1007/s00467-018-4157-2.

15. Pearce D, Manis AD, Nesterov V, Korbmacher C. Regulation of distal tubule sodium transport: mechanisms and roles in homeostasis and pathophysiology. Pflugers Arch - Eur J Physiol 474: 869–884, 2022. doi: 10.1007/s00424-022-02732-5.

16. Rossi GM, Regolisti G, Peyronel F, Fiaccadori E. Recent insights into sodium and potassium handling by the aldosterone-sensitive distal nephron: a review of the relevant physiology. J Nephrol 33: 431–445, 2020. doi: 10.1007/s40620-019-00684-1.

17. Feraille E, Sassi A, Olivier V, Arnoux G, Martin P-Y. Renal water transport in health and disease. Pflugers Arch - Eur J Physiol 474: 841–852, 2022. doi: 10.1007/s00424-022-02712-9.

18. Braakman I, Bulleid NJ. Protein folding and modification in the mammalian endoplasmic reticulum. Annu Rev Biochem 80: 71–99, 2011. doi: 10.1146/annurev-biochem-062209-093836.

19. Schröder M, Kaufman RJ. ER stress and the unfolded protein response. Mutation Research/Fundamental and Molecular Mechanisms of Mutagenesis 569: 29–63, 2005. doi: 10.1016/j.mrfmmm.2004.06.056.

20. Porter AW, Nguyen DN, Clayton DR, Ruiz WG, Mutchler SM, Ray EC, Marciszyn AL, Nkashama LJ, Subramanya AR, Gingras S, Kleyman TR, Apodaca G, Hendershot LM, Brodsky JL, Buck TM. The molecular chaperone GRP170 protects against ER stress and acute kidney injury in mice. JCI Insight 7: e151869, 2022. doi: 10.1172/jci.insight.151869.

21. Buck TM, Kolb AR, Boyd CR, Kleyman TR, Brodsky JL. The Endoplasmic Reticulum–associated Degradation of the Epithelial Sodium Channel Requires a Unique Complement of Molecular Chaperones. Mol Biol Cell 21: 1047–1058, 2010. doi: 10.1091/mbc.E09-11-0944.

22. Buck TM, Plavchak L, Roy A, Donnelly BF, Kashlan OB, Kleyman TR, Subramanya AR, Brodsky JL. The Lhs1/GRP170 Chaperones Facilitate the Endoplasmic Reticulum-associated Degradation of the Epithelial Sodium Channel. J Biol Chem 288: 18366–18380, 2013. doi: 10.1074/jbc.M113.469882.

23. Buck TM, Jordahl AS, Yates ME, Preston GM, Cook E, Kleyman TR, Brodsky JL. Interactions between intersubunit transmembrane domains regulate the chaperone-dependent degradation of an oligomeric membrane protein. Biochem J 474: 357–376, 2017. doi: 10.1042/BCJ20160760.

24. Park J, Easton DP, Chen X, MacDonald IJ, Wang X-Y, Subjeck JR. The Chaperoning Properties of Mouse Grp170, a Member of the Third Family of Hsp70 Related Proteins. Biochemistry 42: 14893–14902, 2003. doi: 10.1021/bi030122e.

25. Schmidt BZ, Perlmutter DH. Grp78, Grp94, and Grp170 interact with α1-antitrypsin mutants that are retained in the endoplasmic reticulum. American Journal of Physiology-Gastrointestinal and Liver Physiology 289: G444–G455, 2005. doi: 10.1152/ajpgi.00237.2004.

26. Cunningham CN, He K, Arunagiri A, Paton AW, Paton JC, Arvan P, Tsai B. Chaperone-Driven Degradation of a Misfolded Proinsulin Mutant in Parallel With Restoration of Wild-Type Insulin Secretion. Diabetes 66: 741–753, 2017. doi: 10.2337/db16-1338.

27. Wang H, Pezeshki AM, Yu X, Guo C, Subjeck JR, Wang X-Y. The Endoplasmic Reticulum Chaperone GRP170: From Immunobiology to Cancer Therapeutics. Front Oncol 4, 2015. doi: 10.3389/fonc.2014.00377.

28. Williams JM, Inoue T, Chen G, Tsai B. The nucleotide exchange factors Grp170 and Sil1 induce cholera toxin release from BiP to enable retrotranslocation. Mol Biol Cell 26: 2181–2189, 2015. doi: 10.1091/mbc.E15-01-0014.

29. Behnke J, Mann MJ, Scruggs F-L, Feige MJ, Hendershot LM. Members of the Hsp70 family recognize distinct types of sequences to execute ER quality control. Mol Cell 63: 739–752, 2016. doi: 10.1016/j.molcel.2016.07.012.

30. Behnke J, Hendershot LM. The Large Hsp70 Grp170 Binds to Unfolded Protein Substrates in Vivo with a Regulation Distinct from Conventional Hsp70s. J Biol Chem 289: 2899–2907, 2014. doi: 10.1074/jbc.M113.507491.

31. Kusaczuk M, Cechowska-Pasko M. Molecular Chaperone ORP150 in ER Stress–related Diseases. Current Pharmaceutical Design 19: 2807–2818, 2013.

32. Bando Y, Tsukamoto Y, Katayama T, Ozawa K, Kitao Y, Hori O, Stern DM, Yamauchi A, Ogawa S. ORP150/HSP12A protects renal tubular epithelium from ischemia-induced cell death. FASEB J 18: 1401–1403, 2004. doi: 10.1096/fj.03-1161fje.

33. Ohse T, Inagi R, Tanaka T, Ota T, Miyata T, Kojima I, Ingelfinger JR, Ogawa S, Fujita T, Nangaku M. Albumin induces endoplasmic reticulum stress and apoptosis in renal proximal tubular cells. Kidney International 70: 1447–1455, 2006. doi: 10.1038/sj.ki.5001704.

34. Boscardin E, Perrier R, Sergi C, Maillard M, Loffing J, Loffing-Cueni D, Koesters R, Rossier BC, Hummler E. Severe hyperkalemia is rescued by low-potassium diet in renal βENaC-deficient mice. Pflugers Arch - Eur J Physiol 469: 1387–1399, 2017. doi: 10.1007/s00424-017-1990-2.

35. Perrier R, Boscardin E, Malsure S, Sergi C, Maillard MP, Loffing J, Loffing-Cueni D, Sørensen MV, Koesters R, Rossier BC, Frateschi S, Hummler E. Severe Salt-Losing Syndrome and Hyperkalemia Induced by Adult Nephron-Specific Knockout of the Epithelial Sodium Channel α-Subunit. J Am Soc Nephrol 27: 2309–2318, 2016. doi: 10.1681/ASN.2015020154.

36. Ray EC, Pitzer A, Lam T, Jordahl A, Patel R, Ao M, Marciszyn A, Winfrey A, Barak Y, Sheng S, Kirabo A, Kleyman TR. Salt sensitivity of volume and blood pressure in a mouse with globally reduced ENaC γ-subunit expression. American Journal of Physiology-Renal Physiology 321: F705–F714, 2021. doi: 10.1152/ajprenal.00559.2020.

37. Cao XR, Shi PP, Sigmund RD, Husted RF, Sigmund CD, Williamson RA, Stokes JB, Yang B. Mice heterozygous for β-ENaC deletion have defective potassium excretion. American Journal of Physiology-Renal Physiology 291: F107–F115, 2006. doi: 10.1152/ajprenal.00159.2005.

38. Mann MJ, Melendez-Suchi C, Sukhoplyasova M, Flory AR, Irvine MC, Iyer AR, Vorndran H, Guerriero CJ, Brodsky JL, Hendershot LM, Buck TM. Loss of Grp170 results in catastrophic disruption of endoplasmic reticulum functions. bioRxiv: 2023.10.19.563191, 2023.

39. Inagi R, Ishimoto Y, Nangaku M. Proteostasis in endoplasmic reticulum—new mechanisms in kidney disease. Nature Reviews Nephrology 10: 369–378, 2014. doi: 10.1038/nrneph.2014.67.

40. Cybulsky AV. Endoplasmic reticulum stress, the unfolded protein response and autophagy in kidney diseases. Nature Reviews Nephrology 13: 681–696, 2017. doi: 10.1038/nrneph.2017.129.

41. Travers KJ, Patil CK, Wodicka L, Lockhart DJ, Weissman JS, Walter P. Functional and Genomic Analyses Reveal an Essential Coordination between the Unfolded Protein Response and ER-Associated Degradation. Cell 101: 249–258, 2000. doi: 10.1016/S0092-8674(00)80835-1.

42. Hetz C, Zhang K, Kaufman RJ. Mechanisms, regulation and functions of the unfolded protein response. Nat Rev Mol Cell Biol 21: 421–438, 2020. doi: 10.1038/s41580-020-0250-z.

43. Walter P, Ron D. The Unfolded Protein Response: From Stress Pathway to Homeostatic Regulation. Science 334: 1081–1086, 2011. doi: 10.1126/science.1209038.

44. Chiti F, Kelly JW. Small molecule protein binding to correct cellular folding or stabilize the native state against misfolding and aggregation. Curr Opin Struct Biol 72: 267–278, 2022. doi: 10.1016/j.sbi.2021.11.009.

45. Kusaczuk M. Tauroursodeoxycholate—Bile Acid with Chaperoning Activity: Molecular and Cellular Effects and Therapeutic Perspectives. Cells 8: 1471, 2019. doi: 10.3390/cells8121471.

46. Traykova-Brauch M, Schönig K, Greiner O, Miloud T, Jauch A, Bode M, Felsher DW, Glick AB, Kwiatkowski DJ, Bujard H, Horst J, Von Knebel Doeberitz M, Niggli FK, Kriz W, Gröne HJ, Koesters R. An efficient and versatile system for acute and chronic modulation of renal tubular function in transgenic mice. Nature Medicine 14: 979–984, 2008. doi: 10.1038/nm.1865.

47. Spanuchart I, Watanabe H, Aldan T, Chow D, Ng RCK. Are Salt Tablets Effective in the Treatment of Euvolemic Hyponatremia? South Med J 113: 125–129, 2020. doi: 10.14423/SMJ.0000000000001075.

48. Kellum JA, Romagnani P, Ashuntantang G, Ronco C, Zarbock A, Anders H-J. Acute kidney injury. Nat Rev Dis Primers 7: 52, 2021. doi: 10.1038/s41572-021-00284-z.

49. Molitoris BA, Sandoval RM, Yadav SPS, Wagner MC. Albumin uptake and processing by the proximal tubule: physiological, pathological, and therapeutic implications. Physiol Rev 102: 1625–1667, 2022. doi: 10.1152/physrev.00014.2021.

50. Schrezenmeier EV, Barasch J, Budde K, Westhoff T, Schmidt-Ott KM. Biomarkers in acute kidney injury – pathophysiological basis and clinical performance. Acta Physiol (Oxf) 219: 554–572, 2017. doi: 10.1111/apha.12764.

51. Jana S, Mitra P, Roy S. Proficient Novel Biomarkers Guide Early Detection of Acute Kidney Injury: A Review. Diseases 11: 8, 2023. doi: 10.3390/diseases11010008.

52. Barnett AM, Babcock MC, Watso JC, Migdal KU, Gutiérrez OM, Farquhar WB, Robinson AT. High dietary salt intake increases urinary NGAL excretion and creatinine clearance in healthy young adults. American Journal of Physiology-Renal Physiology 322: F392–F402, 2022. doi: 10.1152/ajprenal.00240.2021.

53. Nourbakhsh N, Singh P. Role of Renal Oxygenation and Mitochondrial Function in the Pathophysiology of Acute Kidney Injury. Nephron Clinical Practice 127: 149–152, 2014. doi: 10.1159/000363545.

54. Haapaniemi EM, Fogarty CL, Keskitalo S, Katayama S, Vihinen H, Ilander M, Mustjoki S, Krjutškov K, Lehto M, Hautala T, Eriksson O, Jokitalo E, Velagapudi V, Varjosalo M, Seppänen M, Kere J. Combined immunodeficiency and hypoglycemia associated with mutations in hypoxia upregulated 1. Journal of Allergy and Clinical Immunology 139: 1391–1393.e11, 2017. doi: 10.1016/j.jaci.2016.09.050.

55. Rao S, Oyang L, Liang J, Yi P, Han Y, Luo X, Xia L, Lin J, Tan S, Hu J, Wang H, Tang L, Pan Q, Tang Y, Zhou Y, Liao Q. Biological Function of HYOU1 in Tumors and Other Diseases. Onco Targets Ther 14: 1727–1735, 2021. doi: 10.2147/OTT.S297332.

56. Kitao Y, Hashimoto K, Matsuyama T, Iso H, Tamatani T, Hori O, Stern DM, Kano M, Ozawa K, Ogawa S. ORP150/HSP12A Regulates Purkinje Cell Survival: A Role for Endoplasmic Reticulum Stress in Cerebellar Development. J Neurosci 24: 1486–1496, 2004. doi: 10.1523/JNEUROSCI.4029-03.2004.

57. Tamatani M, Matsuyama T, Yamaguchi A, Mitsuda N, Tsukamoto Y, Taniguchi M, Che YH, Ozawa K, Hori O, Nishimura H, Yamashita A, Okabe M, Yanagi H, Stern DM, Ogawa S, Tohyama M. ORP150 protects against hypoxia/ischemia-induced neuronal death. Nat Med 7: 317–323, 2001. doi: 10.1038/85463.

58. Aleshin AN, Sawa Y, Kitagawa-Sakakida S, Bando Y, Ono M, Memon IA, Tohyama M, Ogawa S, Matsuda H. 150-kDa oxygen-regulated protein attenuates myocardial ischemia–reperfusion injury in rat heart. Journal of Molecular and Cellular Cardiology 38: 517–525, 2005. doi: 10.1016/j.yjmcc.2005.01.001.

59. Wiseman RL, Mesgarzadeh JS, Hendershot LM. Reshaping endoplasmic reticulum quality control through the unfolded protein response. Molecular Cell 82: 1477–1491, 2022. doi: 10.1016/j.molcel.2022.03.025.

60. Taniguchi M, Yoshida H. Endoplasmic reticulum stress in kidney function and disease. Curr Opin Nephrol Hypertens 24: 345–350, 2015. doi: 10.1097/MNH.0000000000000141.

61. Burgos JI, Morell M, Mariángelo JIE, Vila Petroff M. Hyperosmotic stress promotes endoplasmic reticulum stress-dependent apoptosis in adult rat cardiac myocytes. Apoptosis 24: 785–797, 2019. doi: 10.1007/s10495-019-01558-4.

62. Martinez-Carrasco R, Fini ME. Dynasore Protects Corneal Epithelial Cells Subjected to Hyperosmolar Stress in an In Vitro Model of Dry Eye Epitheliopathy. International Journal of Molecular Sciences 24: 4754, 2023. doi: 10.3390/ijms24054754.

63. Crambert G, Ernandez T, Lamouroux C, Roth I, Dizin E, Martin P-Y, Féraille E, Hasler U. Epithelial sodium channel abundance is decreased by an unfolded protein response induced by hyperosmolality. Physiological Reports 2: e12169, 2014. doi: 10.14814/phy2.12169.

64. Preissler S, Ron D. Early Events in the Endoplasmic Reticulum Unfolded Protein Response. Cold Spring Harb Perspect Biol 11: a033894, 2019. doi: 10.1101/cshperspect.a033894.

65. Stadt MM, Layton AT. Sex and species differences in epithelial transport in rat and mouse kidneys: Modeling and analysis [Online]. Frontiers in Physiology 13, 2022. https://www.frontiersin.org/articles/10.3389/fphys.2022.991705 [21 Nov. 2023].

66. Layton AT, Sullivan JC. Recent advances in sex differences in kidney function. American Journal of Physiology-Renal Physiology 316: F328–F331, 2019. doi: 10.1152/ajprenal.00584.2018.

67. Hodeify R, Megyesi J, Tarcsafalvi A, Mustafa HI, Hti Lar Seng NS, Price PM. Gender differences control the susceptibility to ER stress-induced acute kidney injury. Am J Physiol Renal Physiol 304: F875–F882, 2013. doi: 10.1152/ajprenal.00590.2012.

68. Shu S, Wang H, Zhu J, Liu Z, Yang D, Wu W, Cai J, Chen A, Tang C, Dong Z. Reciprocal regulation between ER stress and autophagy in renal tubular fibrosis and apoptosis. Cell Death Dis 12: 1–13, 2021. doi: 10.1038/s41419-021-04274-7.

69. Hendershot LM, Buck TM, Brodsky JL. The Essential Functions of Molecular Chaperones and Folding Enzymes in Maintaining Endoplasmic Reticulum Homeostasis..

70. Needham PG, Guerriero CJ, Brodsky JL. Chaperoning Endoplasmic Reticulum–Associated Degradation (ERAD) and Protein Conformational Diseases. Cold Spring Harb Perspect Biol 11: a033928, 2019. doi: 10.1101/cshperspect.a033928.

71. Bleich M, Warth R, Schmidt-Hieber M, Schulz-Baldes A, Hasselblatt P, Fisch D, Berger S, Kunzelmann K, Kriz W, Schütz G, Greger R. Rescue of the mineralocorticoid receptor knock-out mouse. Pflügers Arch 438: 245–254, 1999. doi: 10.1007/s004240050906.

72. Bishop JA, Sharma R, Westra WH. PAX8 immunostaining of anaplastic thyroid carcinoma: a reliable means of discerning thyroid origin for undifferentiated tumors of the head and neck. Human Pathology 42: 1873–1877, 2011. doi: 10.1016/j.humpath.2011.02.004.

73. Wu M-Y, Yiang G-T, Liao W-T, Tsai AP-Y, Cheng Y-L, Cheng P-W, Li C-Y, Li C-J. Current Mechanistic Concepts in Ischemia and Reperfusion Injury. Cellular Physiology and Biochemistry 46: 1650–1667, 2018. doi: 10.1159/000489241.

74. Kang Z, Chen F, Wu W, Liu R, Chen T, Xu F. UPRmt and coordinated UPRER in type 2 diabetes. Front Cell Dev Biol 10: 974083, 2022. doi: 10.3389/fcell.2022.974083.

75. Maity S, Komal P, Kumar V, Saxena A, Tungekar A, Chandrasekar V. Impact of ER Stress and ER-Mitochondrial Crosstalk in Huntington’s Disease. Int J Mol Sci 23: 780, 2022. doi: 10.3390/ijms23020780.

76. Hetz C, Chevet E, Oakes SA. Proteostasis control by the unfolded protein response. Nat Cell Biol 17: 829–838, 2015. doi: 10.1038/ncb3184.

77. Behnke J, Feige MJ, Hendershot LM. BiP and its Nucleotide Exchange Factors Grp170 and Sil1: Mechanisms of Action and Biological Functions. J Mol Biol 427: 1589–1608, 2015. doi: 10.1016/j.jmb.2015.02.011.

78. Wang C-T, Tezuka T, Takeda N, Araki K, Arai S, Miyazaki T. High salt exacerbates acute kidney injury by disturbing the activation of CD5L/apoptosis inhibitor of macrophage (AIM) protein. PLoS One 16: e0260449, 2021. doi: 10.1371/journal.pone.0260449.

79. Brocker C, Thompson DC, Vasiliou V. The role of hyperosmotic stress in inflammation and disease. Biomol Concepts 3: 345–364, 2012. doi: 10.1515/bmc-2012-0001.

80. Tooka K, Darya G, Mina H, Hannaneh G. Impact of high salt diets on CHOP-mediated apoptosis and renal fibrosis in a rat model. Molecular Biology Reports 48: 6423–6433, 2021. doi: 10.1007/s11033-021-06644-y.

81. Neugarten J, Golestaneh L, Kolhe NV. Sex differences in acute kidney injury requiring dialysis. BMC Nephrol 19: 131, 2018. doi: 10.1186/s12882-018-0937-y.

82. Schiffl H. Gender differences in the susceptibility of hospital-acquired acute kidney injury: more questions than answers. Int Urol Nephrol 52: 1911–1914, 2020. doi: 10.1007/s11255-020-02526-7.

83. Müller V, Losonczy G, Heemann U, Vannay Á, Fekete A, Reusz G, Tulassay T, Szabó AJ. Sexual dimorphism in renal ischemia-reperfusion injury in rats: Possible role of endothelin. Kidney International 62: 1364–1371, 2002. doi: 10.1111/j.1523-1755.2002.kid590.x.

84. Park KM, Kim JI, Ahn Y, Bonventre AJ, Bonventre JV. Testosterone is responsible for enhanced susceptibility of males to ischemic renal injury. J Biol Chem 279: 52282–52292, 2004. doi: 10.1074/jbc.M407629200.

85. Veiras LC, Girardi ACC, Curry J, Pei L, Ralph DL, Tran A, Castelo-Branco RC, Pastor-Soler N, Arranz CT, Yu ASL, McDonough AA. Sexual Dimorphic Pattern of Renal Transporters and Electrolyte Homeostasis. J Am Soc Nephrol 28: 3504–3517, 2017. doi: 10.1681/ASN.2017030295.

86. Chen L, Chou C-L, Yang C-R, Knepper MA. Multiomics Analyses Reveal Sex Differences in Mouse Renal Proximal Subsegments. Journal of the American Society of Nephrology 34: 829, 2023. doi: 10.1681/ASN.0000000000000089.

