## Supplemental Figures 1-2 for "Excess dietary sodium partially restores salt and water homeostasis caused by loss of the endoplasmic reticulum molecular chaperone, GRP170, in the mouse nephron"

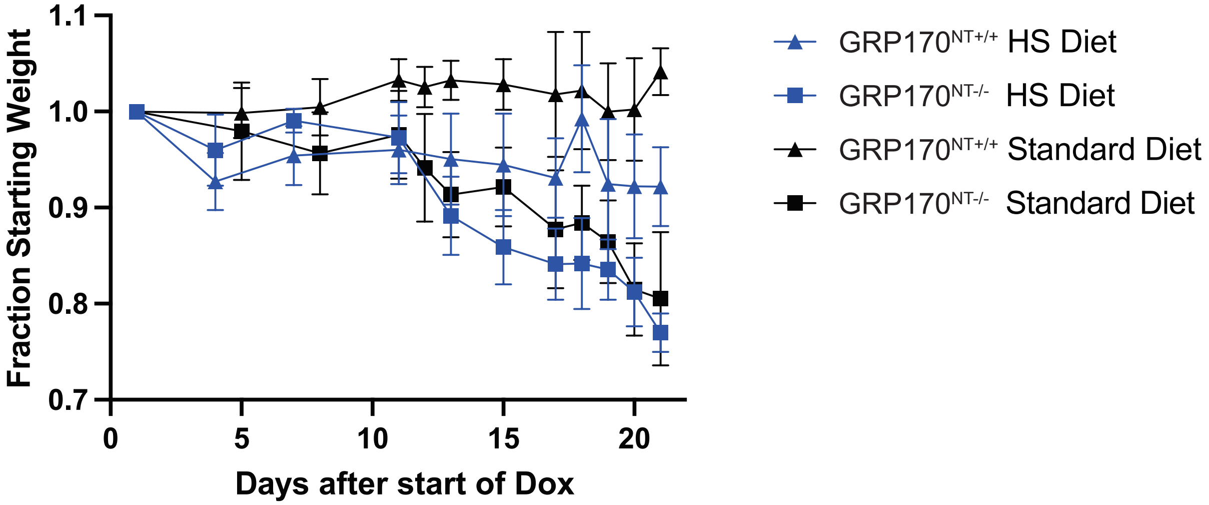


**Supplemental Figure 1: High sodium diet does not prevent weight loss in response to loss of GRP170 in the GRP170$^{NT-/-}$ mice.** Control or GRP170$^{NT-/-}$ mice were treated with dox and weight was monitored a minimum of every third day. Data represent the mean +/-SD; (n=5-12).

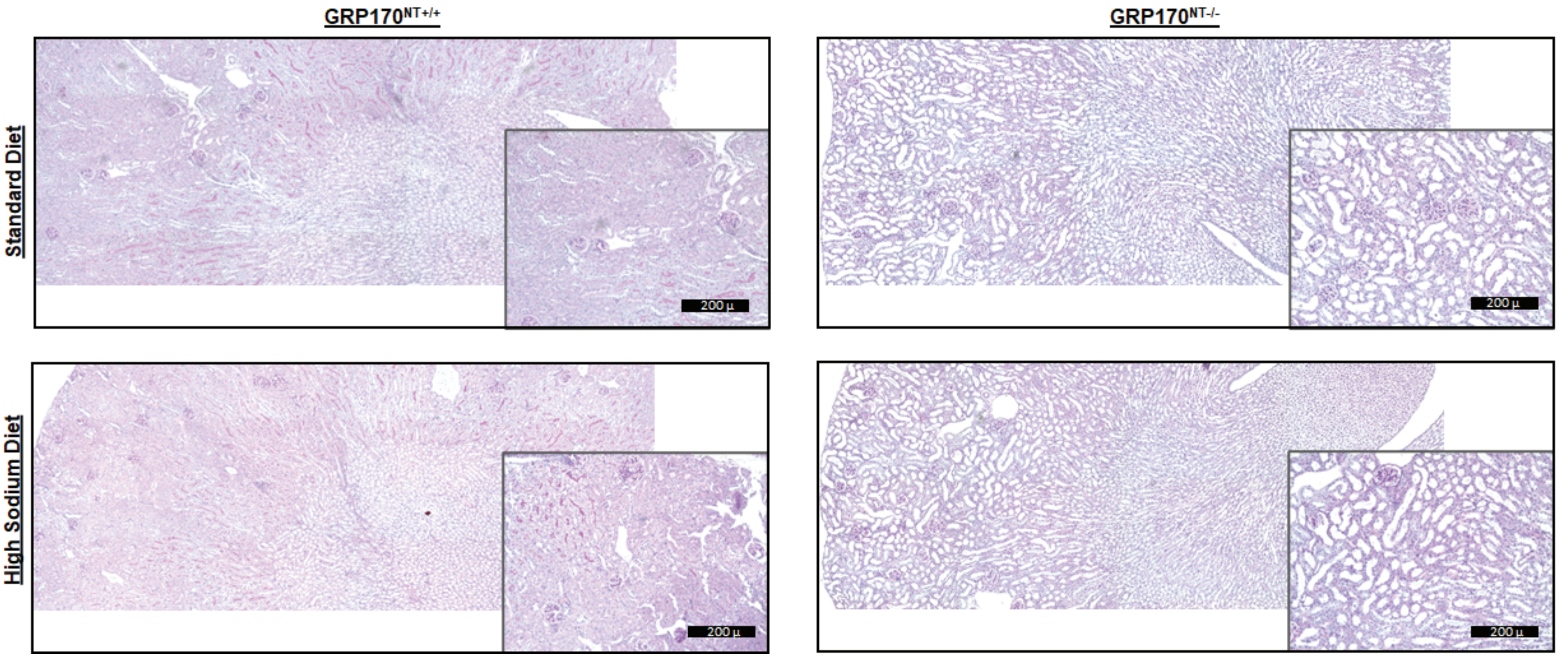


**Supplemental Figure 2: High sodium diet does not prevent kidney injury caused by loss of GRP170 expression in the nephron epithelium of male mice.** Representative PAS staining of kidney sections from male GRP170$^{NT+/+}$ and GRP170$^{NT-/-}$ mice fed eith a standard or an 8% sodium diet are shown. GRP170$^{NT-/-}$ mice exhibit histological findings characteristic of kidney injury: tubular epithelial thinning and dilation, epithelial cell sloughing, and granualar casts. Scale bar: 200 microns.
